# Gradual Warming Drives Life-History Shifts and Reveals Reproductive Ceilings in *Daphnia magna*

**DOI:** 10.1101/2025.08.16.670526

**Authors:** Nikola Petkovic, Tuğçe Ünlü, İsmail K. Sağlam

**Affiliations:** College of Sciences, Molecular Biology and Genetics Department, Koç University, Istanbul, Turkey

**Keywords:** climate change, thermal tolerance, life-history evolution, phenotypic plasticity, transgenerational effects, reproductive thresholds

## Abstract

Global warming often pushes species toward their physiological limits, yet the pace of change may be as important as the absolute temperature reached. Most experiments rely on abrupt shifts, leaving the demographic and evolutionary consequences of gradual warming poorly understood. We conducted a 30-generation experiment with clonal populations of *Daphnia magna* maintained either under constant temperature (26 °C) or gradual warming (+1 °C every 4–6 generations, up to 32 °C). We measured fecundity, growth rate, age at maturation, body size, and lifespan across generations, and used a reciprocal transplant assay to disentangle genetic and plastic responses.

Warming populations matured earlier, grew faster, and initially reproduced more than controls, but suffered shorter lifespans and lost their reproductive advantage near 30 °C, a sub-lethal ceiling that coincided with extinction events. This rate-dependent trajectory contrasts with one-step warming studies that typically report uniformly reduced reproduction. Reciprocal transplants revealed a persistent reduction in adult body size of warming-line offspring, regardless of rearing environment, consistent with heritable or trans-generational change, while growth rate remained largely plastic. Together, these results show that gradual warming can simultaneously elicit plastic and heritable trait shifts, and that reproductive ceilings well below lethal limits act as tipping points for population persistence.

## 1. Introduction

Adverse anthropogenic impacts on the Biosphere have produced environmental change at a rate unprecedented in the Cenozoic (Barnosky et al., 2012). One of the most alarming consequences of the widely recognized issue of global change (Falkowski, 1994; Chivian and Bernstein, 2008; Barnosky et al., 2012) is the shifting of environmental conditions outside of the physiological tolerance range of many species (Parmesan, 2006). Recent studies predict the extinction of 14-35 % of extant species in the following decades (Thomas et al., 2004; Weins and Zelinka, 2024). Hence, understanding the factors which affect adaptation to changing environments has become one of the central problems of contemporary science, with the main goal to help mitigate the detrimental effects of global change. One of the most threatening aspects of global change is the continuous rise of temperature from the pre-industrial era (Hawkins et al., 2017) onwards, a trend predicted to accelerate within the next 50 years (Walther et al., 2002). Consequently, persistence of many extant species may be threatened due to the altered interspecies interactions (for example, food availability) and the limited physiological tolerances to high temperatures (Cahill et al., 2013). The latter factor is particularly pronounced in the lower latitudes, as many tropical species are already experiencing conditions close to their thermal optima (Deutsch et al., 2008). Crucially, population collapse may occur even when lethal limits are not reached, if sub-lethal thresholds for key fitness traits, such as reproduction, are exceeded (Kingsolver and Huey, 2008; Seefeldt and Ebert, 2019).

Predicting the patterns of species’ response to climate change requires tracking of the variation in fitness-related traits (i.e. life histories) during exposure to thermally changing environments, because these traits may strongly correlate with the probability of survival (Isaac, 2009). For example, slower life histories (long gestation and small litters) may enhance extinction risk due to the limited capacity of a population to recover from a decline caused by the natural or anthropogenically induced factors (Purvis et al., 2000). Moreover, these extinction-promoting traits are often negatively correlated with body size (McKinney 1997; Cardillo et al. 2005). Thus, in species with an intraspecific variation in body size, we might expect a shift to a smaller body size due to a lower extinction risk (Isaac, 2009).

However, the opposite pattern can be predicted if an intraspecific variation in body size positively correlates with competitiveness (Isaac, 2009), survival (Yoccoz & Mesnager 1998) and fecundity (Du et al. 2005).

Given the complex interactions among life history traits, clarifying their relative contributions to fitness and survival under rising temperatures requires experimental studies that manipulate temperature and directly monitor trait changes. To date, most of these studies have been carried out using insects as model organisms or various plankton species cultivated in the outdoor mesocosms (Van Doorslaer et al., 2007; Van Doorslaer et al., 2010; Musolin et al., 2010; Ciota et al., 2014; Geerts et al., 2015; Laughton et al., 2017; Cambronero et al., 2018; Iltis et. al., 2019; de Leon et al., 2023; Albini et al., 2025).

Most experimental studies with insects revealed negative correlation of temperature with lifespan, development time, and body size (e.g. Ciota et al., 2014; Laughton et al., 2017; Iltis et. al., 2019). Importantly, these effects of an elevated temperature on the life history traits may depend on the different phenological parameters such as season. For example, an increase in temperature during the warmest part of the season was found to reduce development and increase mortality in the heteropteran species *Nezara viridula*, while adults showed a decline in body size as well as reduced life span and fecundity (Musolin et al., 2010). The negative effects of elevated temperature on life history traits may also vary with geographical distribution of species. In a study of four *Protaphorura* species (Collembola) sampled across a latitudinal gradient, heat stress reduced fecundity in all species, but the most severe effects were observed in the boreal species, which was the most heat-sensitive (de Leon et al., 2023).

Another common experimental approach includes monitoring of freshwater populations under semi-natural conditions (outdoor mesocosms). Mean body size of the plankton species showed variation in response to the elevated temperatures, depending on their cultivation environment. A consistent decline in mean body size has been observed under temperature increases of up to 8°C compared to ambient conditions, including in both single-species setups (Cambronero et al., 2018) and more complex multispecies zooplankton communities (Albini et al., 2025). However, increases in body size at maturity have also been reported under moderate warming (+4°C), likely due to interactions between abiotic factors (e.g., temperature) and biotic factors such as interspecific competition (Van Doorslaer et al., 2010).

In addition, evidence points to latitude-specific differences in how elevated temperature affects fecundity, with contrasting patterns reported for boreal and temperate species. Whereas boreal species typically show reduced fecundity at elevated temperatures (Giovannini et al., 2023), the pattern in temperate-origin species is less consistent, having been reported as positive (Van Doorslaer et al., 2007), negative (Cambronero et al., 2018) or more complex (Wang et al., 2023). In contrast, the response of age at maturation to elevated temperatures appears to be relatively uniform regardless of the environmental conditions, with experimental studies repeatedly reporting earlier maturation at higher temperatures (Van Doorslaer et al., 2007; Cambronero et al., 2018; Giovannini et al., 2023; Wang et al., 2023).

Despite substantial experimental research, significant gaps remain in our understanding of how life history traits respond to temperature change, particularly because most studies have focused on abrupt shifts in temperature, typically involving a single-step increase. In reality, many natural populations experience gradual warming, and the pace of change may be as critical to outcomes as the absolute temperatures reached (Killeen et al., 2017). To better reflect ecological reality, we need experimental designs that examine the effects of gradually and continuously changing thermal environments over longer time scales. This is important for two reasons. First, the rate and therefore the impact of global warming varies across regions, showing positive correlation with both latitude (Root et al., 2003; Deutsch et al., 2008) and altitude (Pepin et al., 2022). Second, experimental studies have shown that the adaptive dynamics of populations to stressful environments are strongly influenced by the rate of environmental change (Collins and de Meaux, 2009; Bell and Gonzalez, 2011; Lindsey et al., 2013; Killeen et al., 2017; Petkovic and Colegrave, 2023). This means that warming regimes of identical endpoints may produce qualitatively different demographic and evolutionary trajectories depending on whether the change is abrupt or gradual (Collins and de Meaux, 2009; Lindsey et al., 2013).

Experimental studies with microbial systems highlight the importance of warming rate for adaptive dynamics. For example, populations exposed to the most gradual temperature increases are more likely to persist and evolve higher thermal tolerance, consistent with predictions of evolutionary rescue (Killeen et al., 2017). Similarly, slower environmental change can promote growth at high temperatures but may come at a cost to long-term survival due to correlated selection (Liukkonen et al., 2021). Such findings suggest that gradual warming can initially enhance certain performance traits yet may eventually drive collapse once sub-lethal thresholds for reproduction or other key functions are exceeded (Huey and Kingsolver, 2019). However, such studies have primarily focused on growth rate in unicellular organisms. We still lack a clear understanding of how more complex life history traits, such as body size, fecundity, and age at maturation, respond to gradual thermal change, particularly in multicellular species. Most experimental work on these traits has been based on abrupt or one-step temperature increases, limiting ecological relevance.

In addition, the extent to which these trait changes result from phenotypic plasticity versus genetic adaptation remains poorly understood. Organisms may respond to thermal shifts through adaptive plasticity, adjusting trait expression without genetic change (Lalejini et al., 2021). For example, *Daphnia magna* populations can modify hemoglobin expression in response to temperature (Yampolsky et al., 2014). Yet, genetic responses to warming have also been observed in *D. magna*, including short-term adaptation in laboratory populations (Seefeldt and Ebert, 2019) and long-term responses in the wild (Geerts et al., 2015).

Importantly, plasticity itself can evolve (Schaum et al., 2022), making it difficult to disentangle the roles of plasticity and genetic change in trait evolution. Properly distinguishing between these mechanisms requires experiments that track life history traits under common garden conditions. To date, studies that have attempted such distinctions have almost exclusively focused on abrupt temperature shifts (for example, Van Doorslaer et al., 2010), leaving the effects of gradual temperature deterioration largely unexplored.

To address these gaps, we conducted a controlled multigenerational evolutionary experiment with three clonal lines of *D. magna* to test how multiple life-history traits respond to gradually increasing temperatures, a scenario more representative of ongoing climate change than abrupt thermal shifts. Populations were propagated through non-overlapping generations under either constant conditions (control) or a warming regime that increased by 1 °C every 4–6 generations. We monitored fecundity, somatic growth rate, age at maturation, body size, and extinction risk across generations, and used a reciprocal transplant experiment to disentangle genetic adaptation from phenotypic plasticity. This design allowed us to evaluate both the pace and nature of trait change under sustained warming. Our results provide insight into how gradual thermal deterioration can alter life-history strategies in a key model species, informing predictions of population responses to ongoing climate change. In doing so, our study also identifies a potential tipping point in reproductive performance that precedes mortality, providing a novel perspective on how gradual warming can restructure life-history strategies well before lethal limits are reached.

### 2. Materials and Methods

### 2.1 Experimental procedures

#### Model Organism and Culture Conditions

*Daphnia magna* Straus (1820) is a planktonic freshwater crustacean (class Phyllopoda) that reproduces via cyclic parthenogenesis, producing clutches of up to 100 eggs after each adult molt (Ebert, 2005). Eggs develop directly into juveniles (neonates), bypassing a resting stage. The generation time is temperature-dependent, with the first clutch typically produced 5–10 days after birth at the optimal temperature of 20°C (Ebert, 2005). Due to its clonal reproduction and tractable life cycle, *D. magna* is a widely used model for studying phenotypic plasticity and local adaptation (Yampolsky et al., 2014).

Three clonal lines were used: HU-HO-2 (hereafter referred to as Hu) and TR-EG-1 (hereafter referred to as Tr) (University of Basel), and An (Middle East Technical University, Ankara). Cultures were maintained in 250 ml glass beakers with 150 ml of ADaM medium (Klüttgen et al., 1994), replenished weekly. All populations were kept under a 16:8 h light:dark cycle and fed with *Chlamydomonas reinhardtii* at 50,000 cells/ml three times per week. Temperature treatments varied according to experimental design.

#### Preliminary Thermal Performance Assay

Before the main experiment, we conducted a baseline thermal tolerance assay following Seefeldt and Ebert (2019) to evaluate the performance limits of the three *D. magna* clones and to inform temperature treatment design.

Five replicate populations per clone (Hu, Tr and An) containing five adults were maintained at 22°C in 100 ml beakers containing ADaM medium, replenished weekly. Populations were acclimated to increasing thermal conditions by raising the temperature to 25°C in week one and to 28°C in week two. Juvenile females (1–3 days old) were then sampled from each replicate and individually exposed to one of four test temperatures (30°C, 31°C, 32°C, or 33°C) in 60 ml jars. Each temperature was replicated at least three times per clone. Over two weeks, individuals were monitored daily for evidence of reproduction (presence of eggs in brood chambers or neonates in the medium).

#### Experimental Design and Temperature Treatments

A total of 24 experimental populations were established, comprising 12 treatment and 12 control lines (3 clones × 4 replicates each). Each population consisted of 15 individuals per generation. Non-overlapping generations were maintained by transferring the first 15 neonates to a fresh beaker of medium at each generation (see Figure 1 for more details regarding the experimental design and transfer procedures).

**Figure 1 –.**
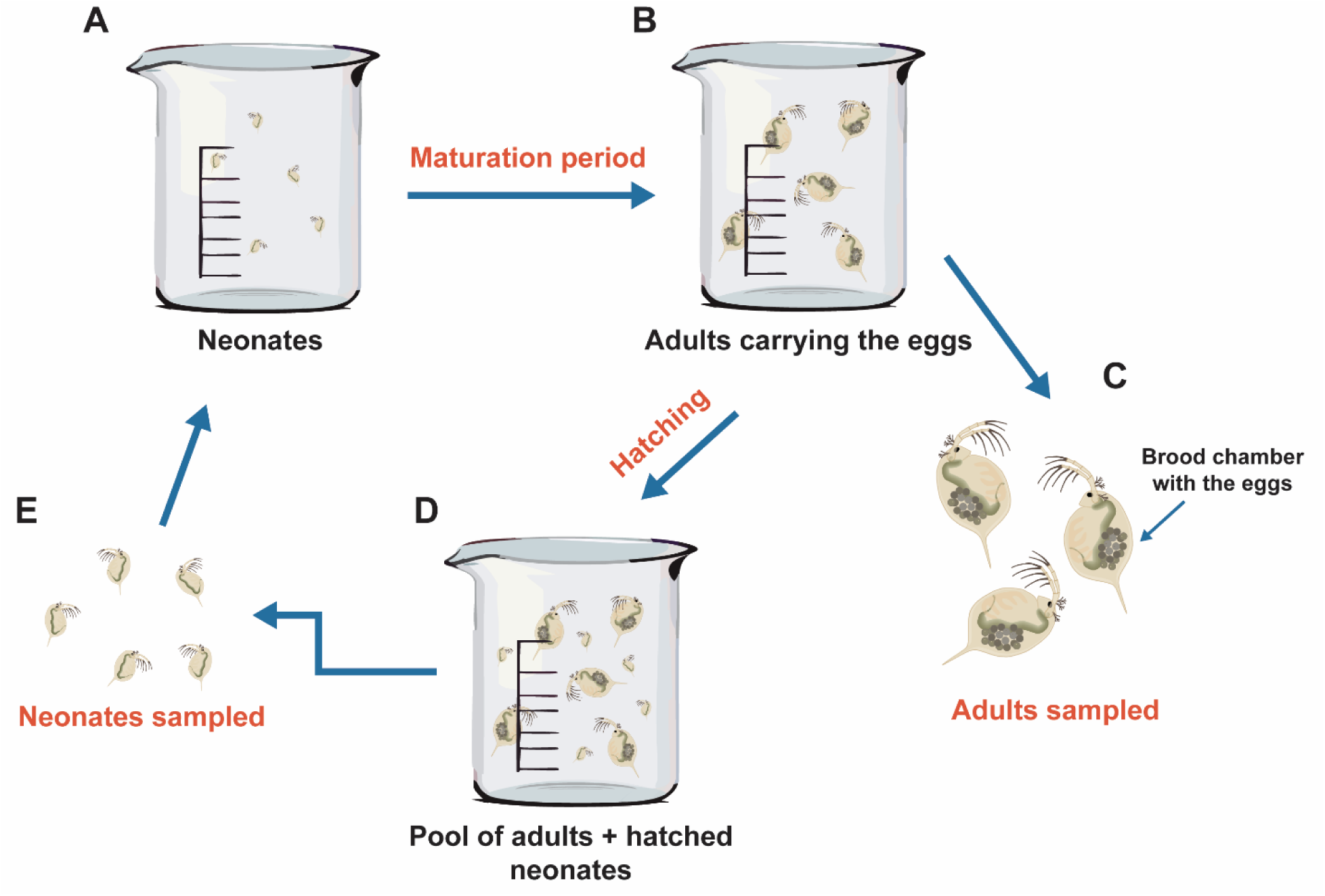
Transfer procedure and data collection regime of experimental populations; (A) 15 neonates per beaker were cultivated until they reached maturation (B), defined as the time when the first eggs appeared in their brood chambers (note that both neonates and adults are downscaled relative to the beakers for an easier representation); C) after each round of temperature increase, 3 adults were sampled for the main assays for quantification of life history traits; D) hatching (the release of the newborn neonates from the brood chambers) resulted in formation of pool of both adults and neonates (note that here, for simplicity, the ratio of newborns vs adults is drawn as 1:1; typically, a larger number of neonates is present); E) after each new generation of neonates appeared, 15 individuals were sampled and transferred to a new beaker (i.e. the next generation of an experimental population has been initiated.

In the treatment group (hereafter referred to as ‘warming’), temperature increased gradually over time, beginning at 26°C. Temperature was raised by 1°C every 4 generations until it reached 29°C, and then by 1°C every 6 generations thereafter. This slowing of the rate beyond 29°C was intended to minimize heat-induced extinction risk, based on prior findings (Mitchell and Lampert, 2000) and the results of our baseline assay.

Control populations were maintained at a constant 26°C for the duration of the experiment.

#### Life History Trait Assays

To quantify evolutionary responses to gradually deteriorating (increasing) temperature, life history traits were measured at five timepoints: 26°C, 27°C, 28°C, 29°C, and 30°C, corresponding to generations 4, 8, 12, 18, and 24 in the treatment populations.

At each timepoint, the first three mature individuals (i.e., individuals with eggs visible in the brood chamber) were isolated from each population and placed individually in 60 ml jars containing ADaM at the population’s current temperature. Each female’s first and second clutches were counted to estimate fecundity. The first clutch was discarded after hatching to avoid size bias, as first-clutch offspring tend to be smaller (Ebert, 1991). After release of the second clutch, the mother was removed.

Six neonates from the second clutch were randomly selected and reared individually in the same jars. Pilot tests had shown that higher densities could trigger unwanted male production or diapause (ephippia) formation. Neonates were photographed at birth and again at maturation using brightfield microscopy (BRESSER Advance ICD Stereo Microscope with Full HD eyepiece camera at 10x magnification). Body size was measured via gut length (GL) of the relaxed animals in ImageJ and converted to dry mass (DM) using the equation: DM = 0.00679 × GL^2.75^ (Fossen et al., 2018).

Neonates were monitored daily for development and survival. Age at maturation was defined as the time when the first eggs were visible in their brood chambers. Lifespan was defined as the number of days from birth until death. Somatic growth rate was calculated as the difference in dry mass between birth and maturation, divided by the number of days between the two measurements. The clutch interval (time between first and second clutches) was also recorded.

#### Extinction Monitoring

Each population was visually inspected daily. Populations were recorded as extinct when no live individuals remained in the beaker.

#### Reciprocal Transplant Assay

To disentangle the effects of phenotypic plasticity and genetic adaptation, a reciprocal transplant was conducted at generation 25. Three mature individuals were sampled from each control and treatment population. From each adult, up to six second-clutch neonates were reared in the environment opposite to their lineage’s ancestral condition.

Control-derived neonates were raised at 30°C, representing novel heat exposure. Treatment-derived neonates were raised at 26°C, representing a return to ancestral temperature. These individuals were reared until maturation, and their adult body size and somatic growth rate were quantified using the same methods as described above.

### 2.2 Data Analysis

#### 2.2.1 Preliminary Thermal Performance Assay

To assess the baseline thermal tolerance of ancestral clones, we fit a generalized linear mixed model (GLMM) with a binomial error distribution. The presence or absence of offspring (eggs or neonates) was treated as a binary response variable. Clone identity was included as a fixed categorical predictor, and temperature was included as a continuous covariate.

#### 2.2.2 Extinction dynamics

The extinction dynamics of each treatment group were analysed by fitting a Weibull regression model with two fixed factors: ‘treatment’ and ‘clone‘. The interaction between the factors was included in the model. The best fit was obtained using the extreme hazard function.

#### 2.2.3 Life History Trait Assays

The datasets containing life history trait values from the main assays were analyzed using linear mixed models (LMMs). The models included “treatment” and “clone” as categorical fixed effects, “generation” as a continuous fixed effect, and “population” and “individual” as categorical random effects to account for hierarchical structure. Each of the seven life history traits was modeled separately as a continuous response variable. Simplified models excluding interaction terms were favored over the more complex models that included the interactions if AIC values were lower (see Appendix section for details). Given the limited sample sizes within each dataset, model parameters were estimated using restricted maximum likelihood (REML). The potential outliers were identified using Grubb’s test and were excluded from model comparison (see Appendix section for details).

The potential covariation among life history traits in the experimental populations was examined using Pearson’s correlation test, based on pairwise comparisons between traits.

#### 2.2.4 The Reciprocal Transplant Assay

The dataset from the reciprocal transplant assay was analyzed using linear mixed models (LMMs). Fixed effects included “environment” (ancestral or novel), “treatment” (control or thermal deterioration), and “clone” (all categorical). Random effects accounted for hierarchical structure by including “population” and “individual” as categorical variables.

Each of the two monitored life history traits—age at maturation and growth rate—was modeled separately as a continuous response variable. Given the limited number of observations in each dataset, model parameters were estimated using restricted maximum likelihood (REML).

All the analyses were carried out using RStudio (version 2024.12.1; R Core Team, 2017). The packages: ‘gplots’ (v3.1.3; Warnes et al., 2022), ‘ggplot2’ (v3.4.2; Wickham, 2016), ‘plothrix’ (v3.8.2; Lemon, 2006), ‘RColorBrewer’ (v1.1-3; Neuwirth, 2022) were used for visualization of the datasets: baseline thermal tolerance of ancestral clones; life history trait assays; the reciprocal transplant assay. The package ‘plyr’ (v1.8.8; Wickham, 2011) was used for summarizing the data prior to visualization. ‘Survival’ (v3.2-13; Therneau, 2021) and ‘survminer’ (v0.4.9; Kassambara et al., 2021) were used for the analysis and visualization, respectively, of extinction dynamics. For fitting of the mixed models, the packages lme4 (Bates et al., 2014) and lmerTest (Kuznetzova et al., 2017) were used.

## 3. Results

### 3.1 Preliminary thermal performance assay

The reproductive ability of the ancestral clones varied with test temperature (Generalised Linear Mixed Model; df = 1; F1,247 = 11.76; P < 0.001). While the average successful reproduction rate at the temperature range between 30°C and 32°C was 28%, only a single individual (0.01%) successfully reproduced at 33°C (Figure 2A). Clone identity also significantly affected reproduction (Generalised Linear Mixed Model; df = 2; F2,246 = 6.7; P = 0.001) (Figure 2B), with the Tr clone showing 2.66 times higher percentage of successfully reproduced individuals relative to the other two clones combined (z = 3.154; P = 0.002). This assay established that the upper thermal limit for successful reproduction in the experimental populations lies just below 33 °C, with sharply reduced success already evident between 30-32 °C.

**Figure 2 –.**
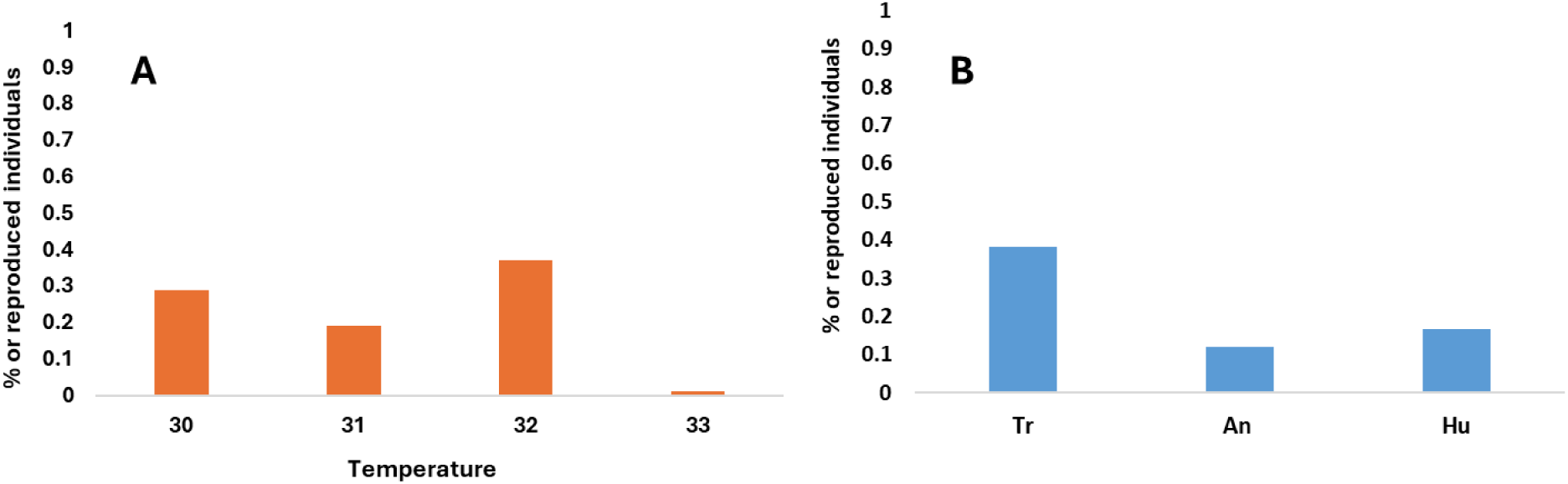
Thermal performance assay of the ancestral clones; **A)** Percentage of successfully reproduced individuals per test temperature; **B)** Percentage of successfully reproduced individuals per clone.

### 3.2 Main evolutionary experiment

#### Population extinction dynamics

Populations in the warming treatment tended to go extinct earlier than controls, although this effect was marginal (Weibull regression; df = 1; χ^2^ = 3.04; P = 0.08) (Figure 3A). Clone identity had a similarly marginal effect on extinction rates (Weibull regression; df = 2; χ^2^ = 5.02; P = 0.08) (Figure 3B).

**Figure 3 –.**
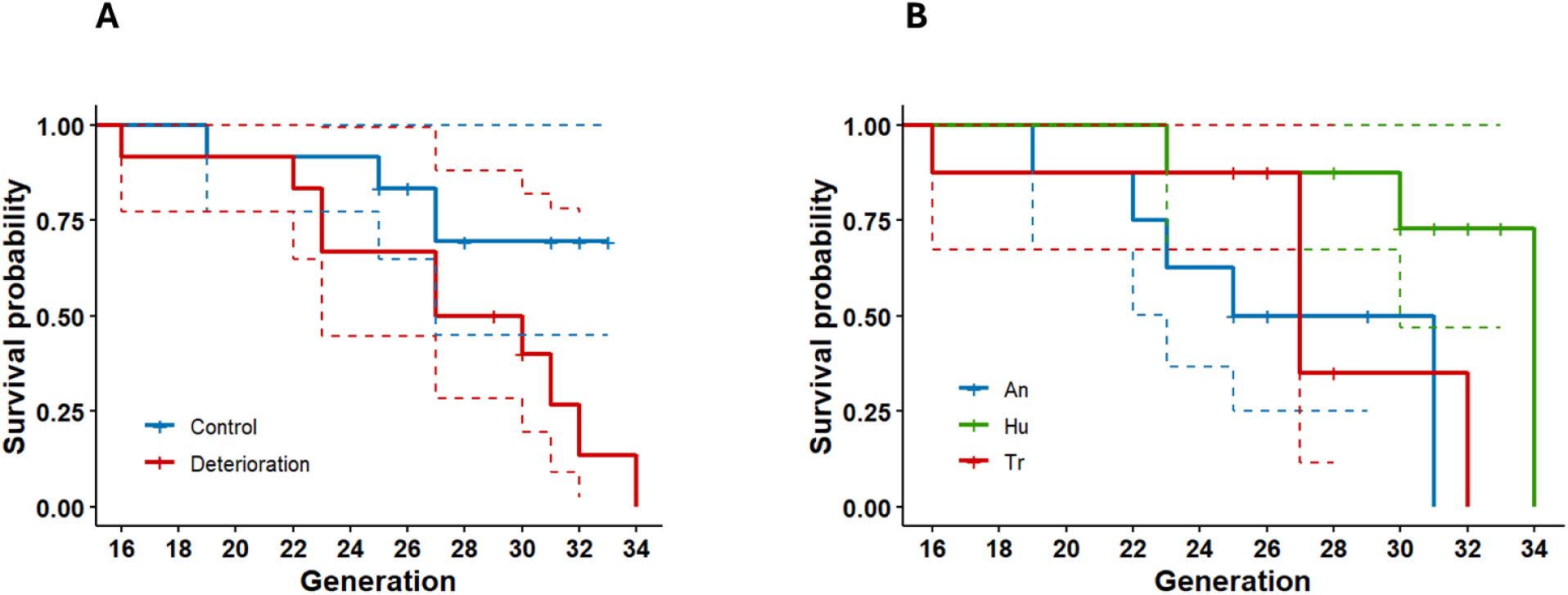
Extinction dynamics of experimental populations: per treatment group. **(A); per clone (B)**; extinctions are plotted against the generations upon each extinction event; 95% confidence interval for each Kaplan-Meier survival curve is represented with the dotted lines (upper and lower boundary).

#### Life-history trait responses

##### Reproductive traits

Fecundity declined significantly across generations in both treatments (LMM, df = 1; F = 41.33; P < 0.001) (Figure 4A) but remained 12% higher in warming populations than controls (LMM, df = 1; F = 8.3; P = 0.004) (Figure 4B). A significant clone-by-treatment interaction (LMM; df = 2; F = 4.06; P = 0.02) was driven by higher fecundity in Hu and Tr clones relative to An under warming (An vs Hu, t = 2.58, P = 0.01; An vs Tr, t = 2.35, P = 0.02; Figure 4D). A generation-by-treatment interaction (LMM; df = 1; F = 4.59; P = 0.03) revealed that this advantage was consistent except at 30 °C (24^th^ generation), where warming-line fecundity was 34% lower than controls (Figure 4C).

**Figure 4 –.**
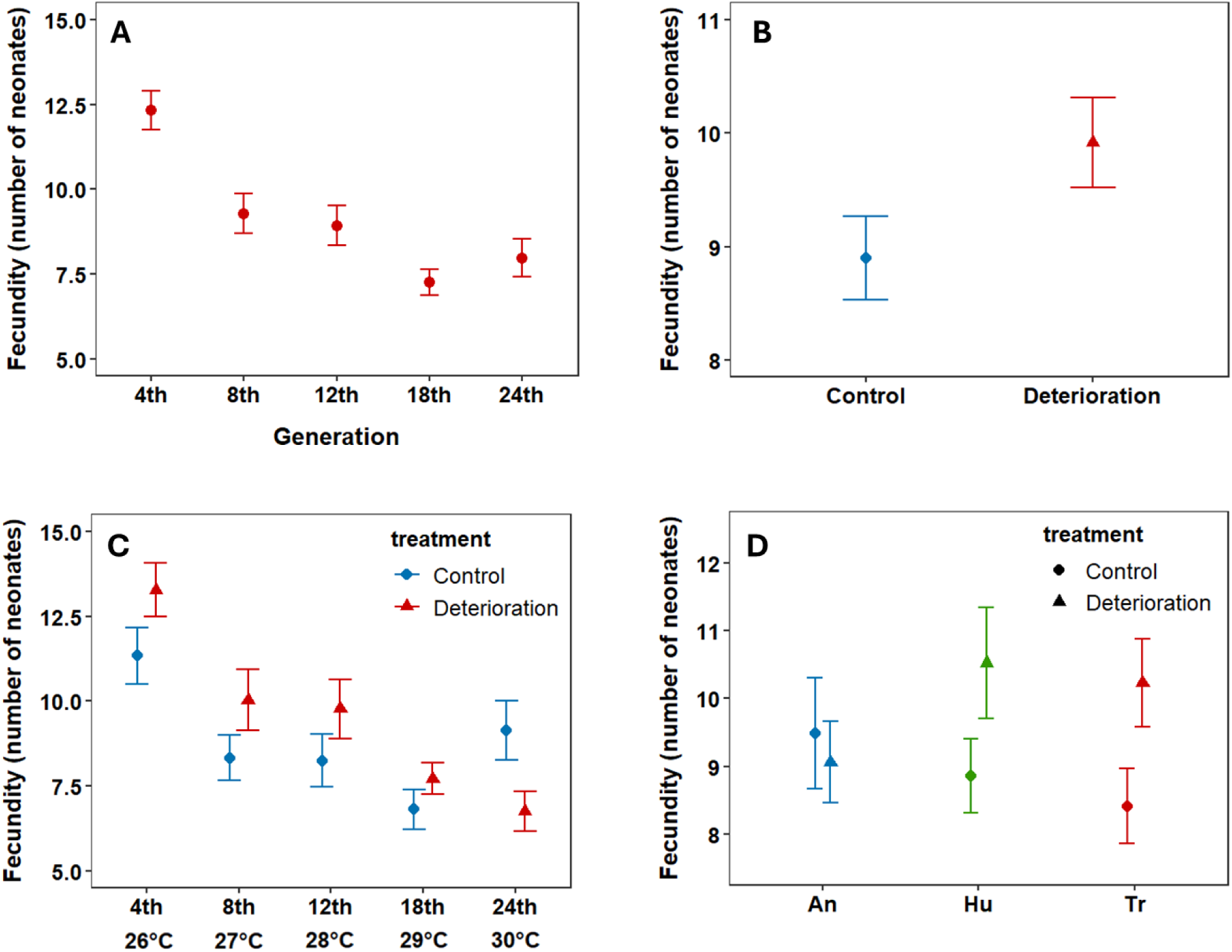
Mean fecundity (the number of neonates within the first two clutches) of the populations: per generation (A); per treatment group (B); per treatment group during each assay (C): X axis contains the number of generations that each assayed population underwent, with the temperature degree that the warming populations experienced during the assay stated below; per clone, per treatment group (D); the bars represent standard error of the mean.

We saw no overall change in span between clutches (LMM, df = 1; F = 1.33; P = 0.25), but a marginal clone-by-treatment interaction (LMM, df = 2; F = 2.96; P = 0.05) indicated shorter intervals for Tr under warming (t = 2.29, P =0.02) (Figure 5A).

**Figure 5 –.**
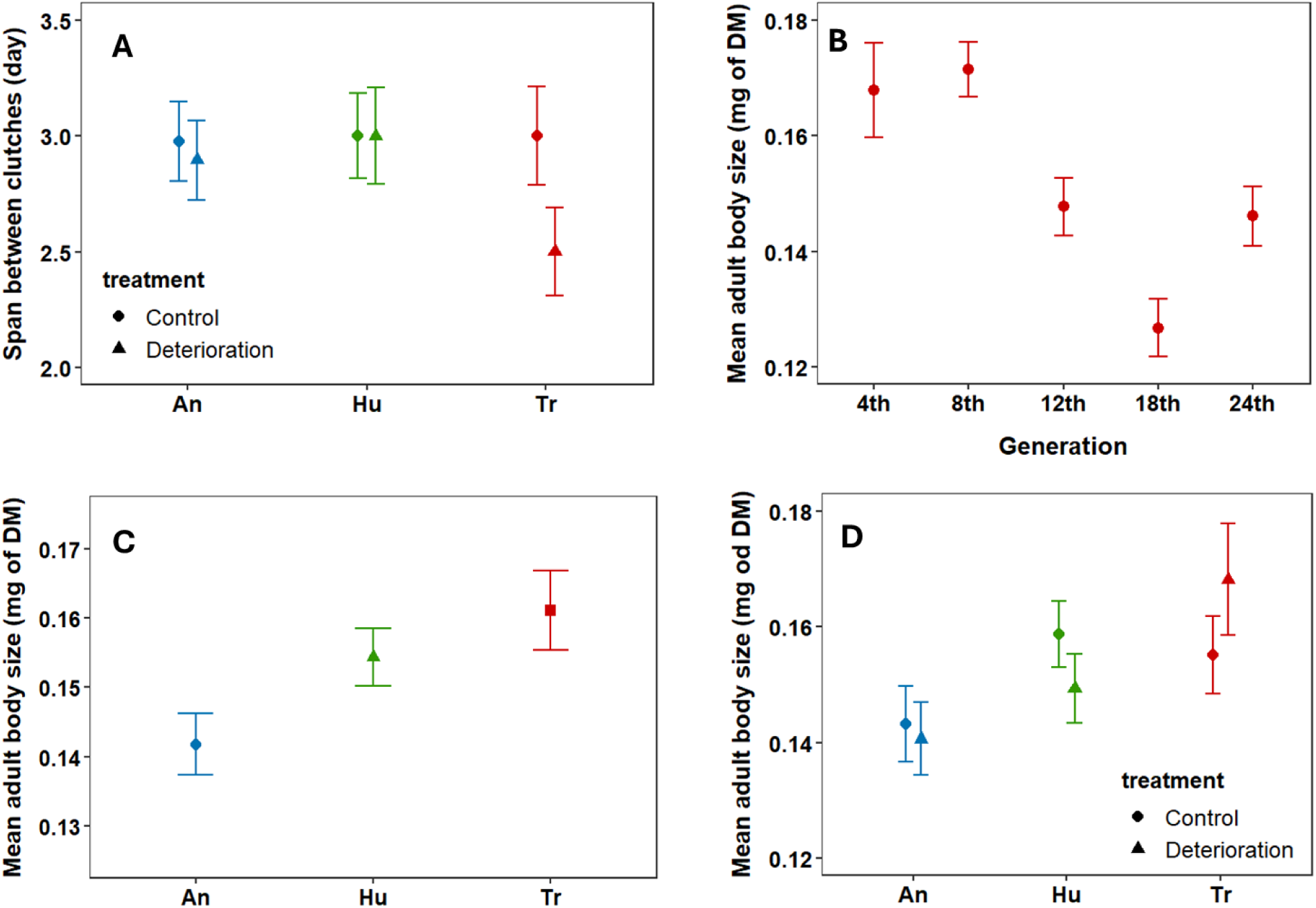
Mean span between the first two clutches (day) of the populations (A); Mean adult body size (mg od DM) of the populations (B-D): per generation (B); per clone (C); per clone per treatment group; the bars represent standard error of the mean.

##### Size traits

Neonate body size was unaffected by generation (LMM, df = 1; F = 1.27; P = 0.26), treatment (LMM, df = 1; F = 1.84; P = 0.18) or clone (LMM, df = 2; F = 1.22; P = 0.3). In contrast, adult body size declined significantly over generations (LMM, df = 1; F = 17.63; P < 0.001) (Figure 5B). Treatment had no main effect (LMM, df = 1; F = 1.48; P = 0.23) but clone identity was significant (LMM, df = 2; F = 3.37; P = 0.04) with mean body size of An declining more than Hu or Tr (Figure 5C). A clone-by-treatment interaction (LMM, df = 2; F = 3.12; P = 0.047) revealed that Tr maintained a size advantage in the warming treatment (Tr vs Hu, t = 2.29, P = 0.02; Tr vs An, t = 1.99, P = 0.048) (Figure 5D).

##### Development and Somatic growth rate

Development time increased with generation (LMM, df = 1; F = 12.56; P = 0.005) (Figure 6A) but was on average 3 days shorter (23% faster maturation) in warming populations (LMM, df = 1; F = 7.66; P = 0.006) (Figure 6B). Clone effects were marginal (LMM, df = 2; F = 2.93; P = 0.05), with Tr maturing later than An (t = 2.41, P = 0.02) (Figure 6C).

**Figure 6 –.**
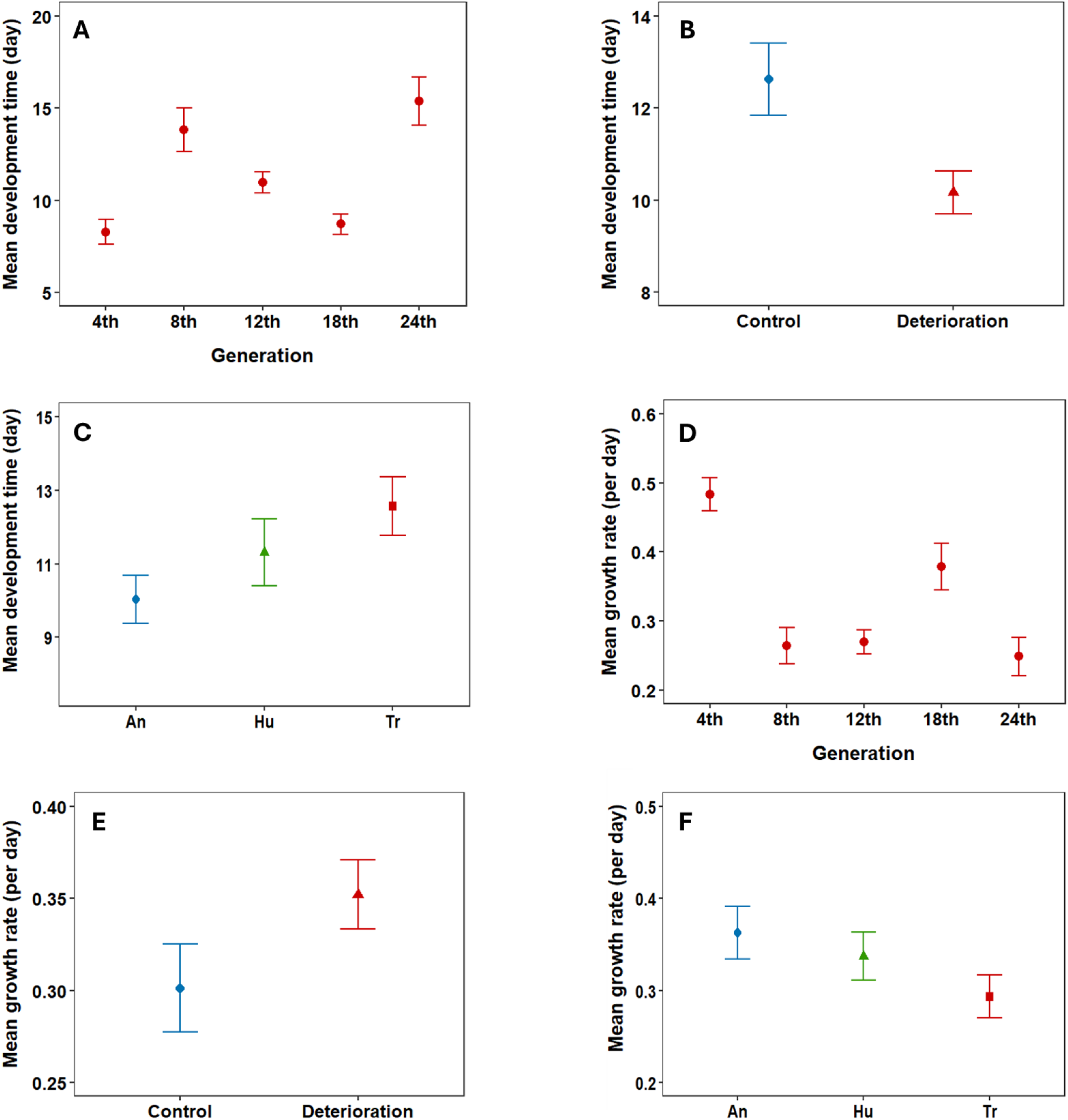
Mean development time (day) of the populations. **(A-C)**: per generation (A); per treatment group (B); per clone (C); **Mean somatic growth rate (per day) of the population (D - F)**: per generation (D); per treatment group (E); per clone (F); the bars represent standard error of the mean.

Somatic growth rate declined overall (LMM, df = 1; F = 23.05; P < 0.001) (Figure 6D) but was 14% faster in warming populations (LMM, df = 1; F = 6.23; P = 0.01) (Figure 6E). Clone effects were marginal (LMM, df = 2; F = 2.43; P = 0.09), with An growing faster than Tr (t = 2.14, P =0.03) (Figure 6F).

##### Lifespan

Lifespan decreased significantly across generations (LMM, df = 1; F = 35.59; P < 0.001) (Figure 7A) and was 21% shorter in warming populations than controls (LMM, df = 1; F = 24.92; P < 0.001) (Figure 7B). The largest treatment effect occurred at 26°C (4^th^ generation) (LMM, generation × treatment; df = 1; F = 10.21; P = 0.002; Figure 7C). Clone identity was significant (LMM, df = 2; F = 7.43; P < 0.001), with An living longer than both Tr (t = - 3.83; P < 0.001) and Hu (t = - 3.16; P = 0.002) (Figure 7D).

**Figure 7 –.**
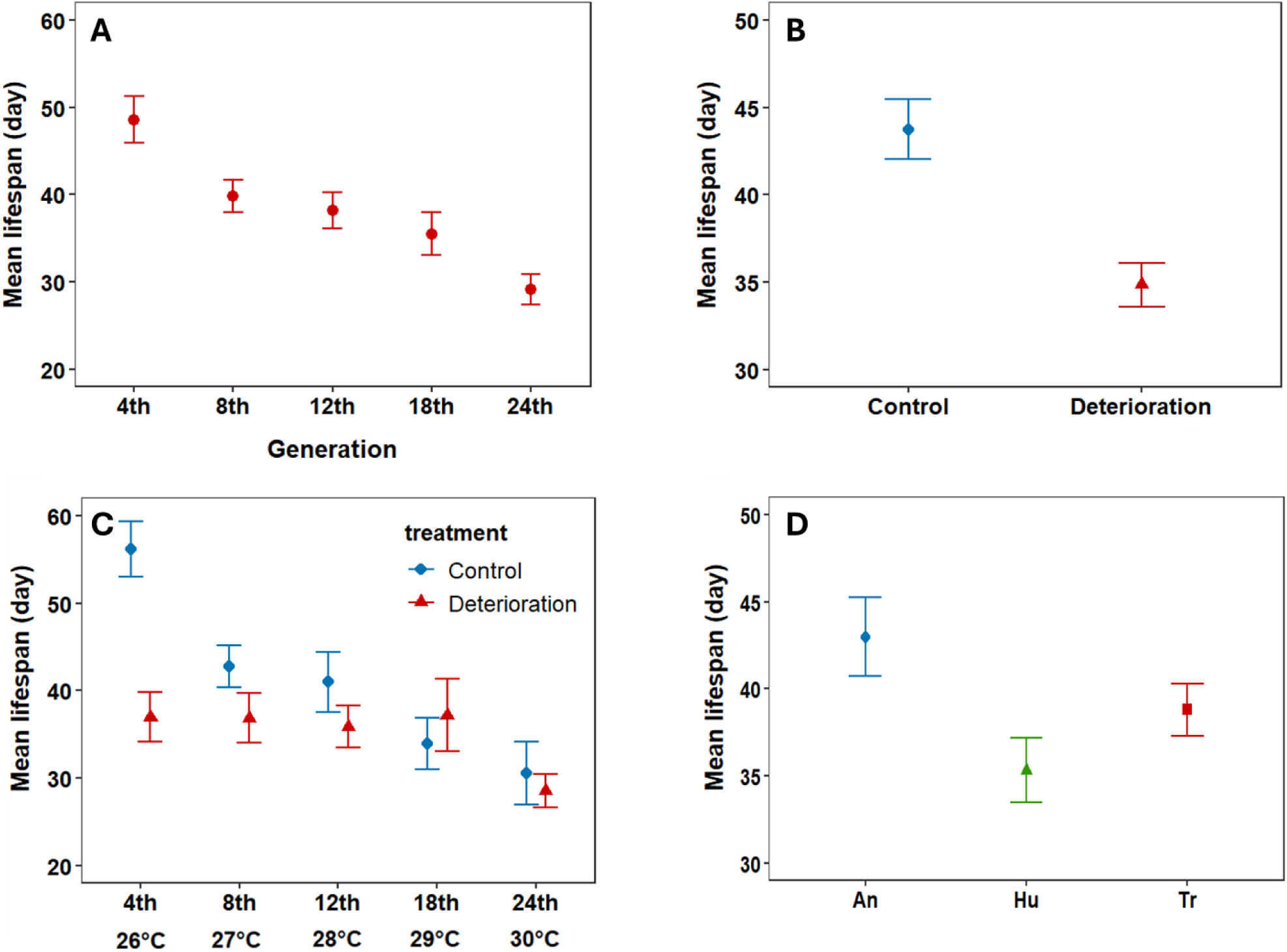
Mean lifespan (day) of experimental populations: per generation (A); per treatment group (B); per treatment group during each assay (C): X axis contains the number of generations that each assayed population underwent, with the temperature degree that the warming populations experienced during the assay stated below; per clone (D); the bars represent standard error of the mean.

##### Life history trait covariation in warming populations

Somatic growth rate was strongly negatively correlated with development time (r = - 0.86; t (60) = - 13.41; P < 0.001) (Figure 8A) and with both neonate body size (r = - 0.31; t (60) = - 2.52; P = 0.01), and adult body size (r = - 0.49; t (60) = - 4.38; P < 0.001) (Figure 8B). Development time was positively correlated with adult body size (r = 0.64; t (60) = 6.42; P < 0.001) (Figure 8C), and adult body size correlated positively with fecundity (r = 0.43; t (67) = 3.92; P = 0.002) (Figure 8D). Other correlations were weak and non-significant (see Appendix section for details).

**Figure 8 –.**
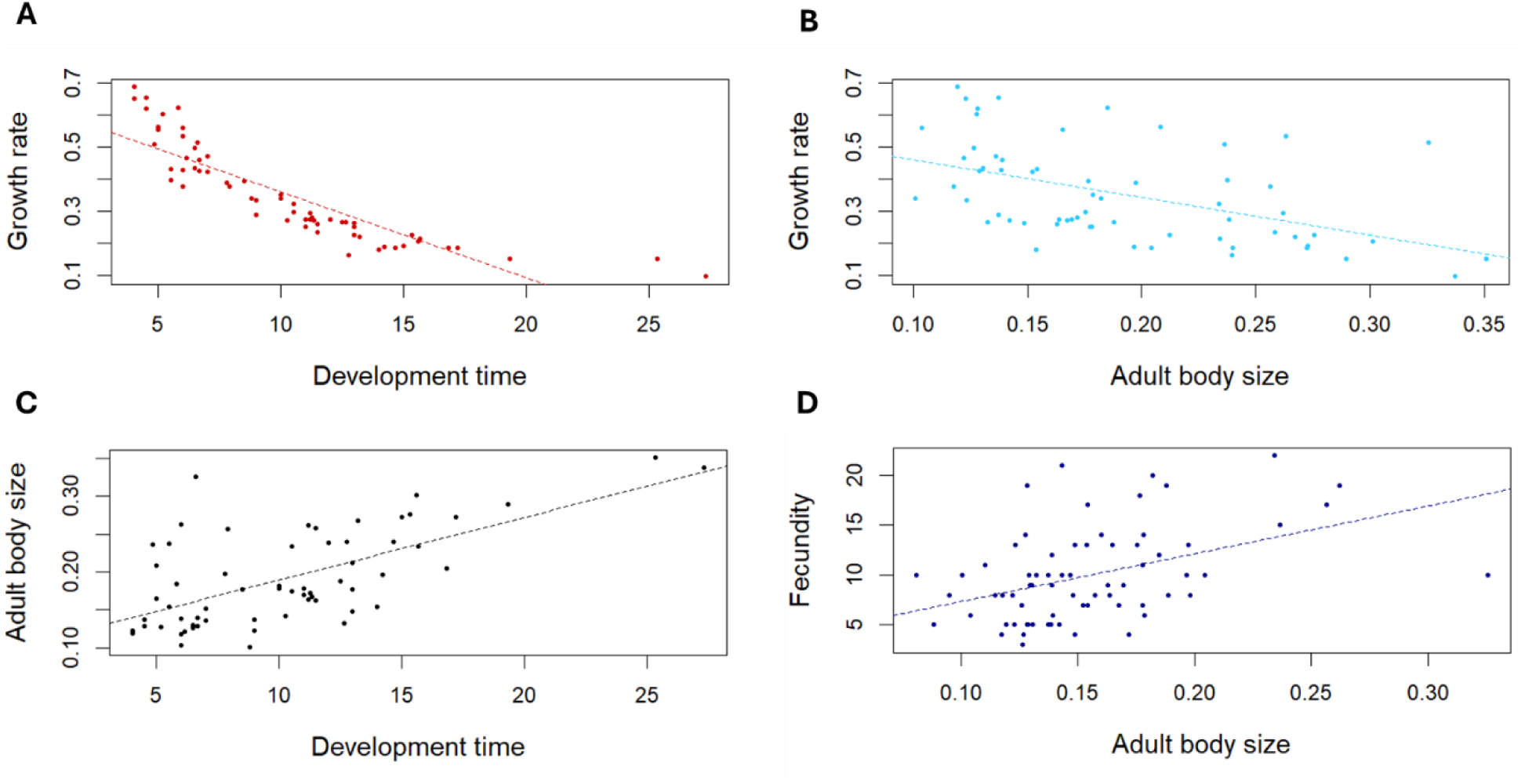
Significant correlations between the pairs of the life history traits in warming populations; **A)** Somatic growth rate (per day) plotted against development time (day); **B)** Somatic growth rate (per day) plotted against adult body size (mg of DM); **C)** Adult body size (mg of DM) plotted against development time (day); **D)** Fecundity (number of neonates within the first two clutches) plotted against adult body size (mg of DM).

### 4.3 The reciprocal transplant experiment

Mean adult body size differed significantly between treatments (LMM, df = 1; F = 9.05; P = 0.004), with control-line offspring 17% larger than warming-line offspring, regardless of rearing environment (LMM, environment × treatment; df = 1; F = 0.32; P = 0.57) (Figure 9A).

**Figure 9 –.**
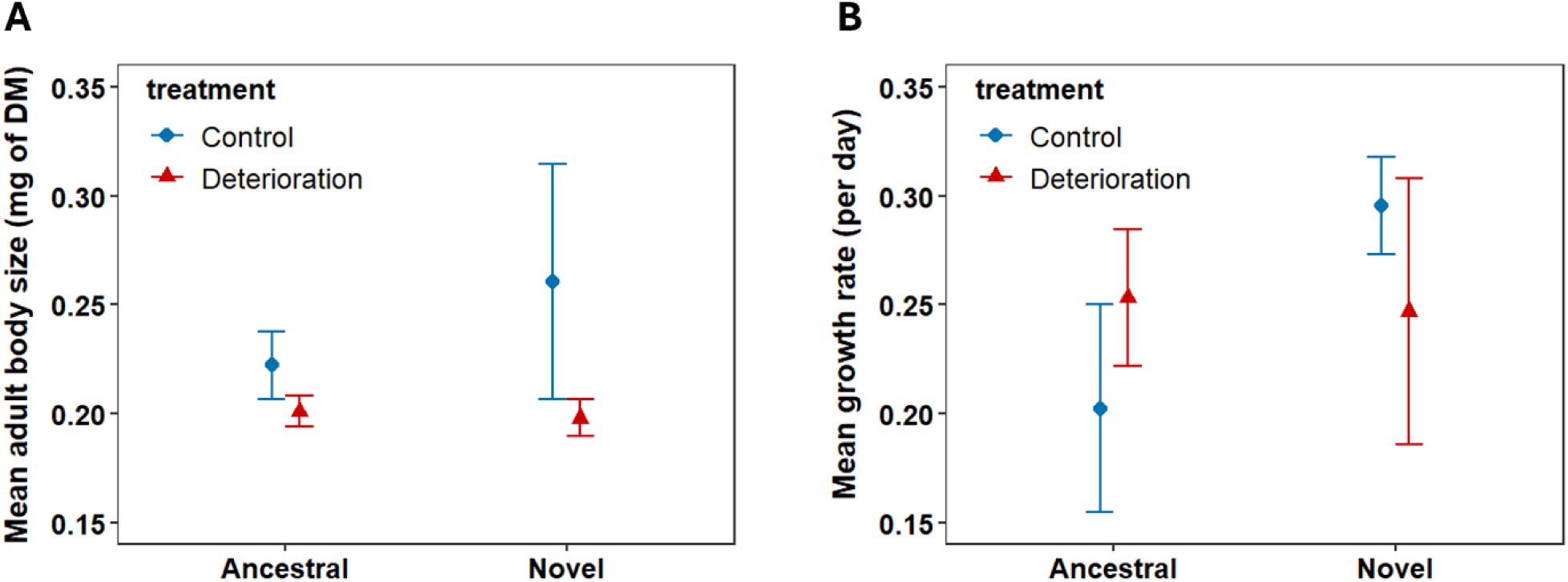
The reciprocal transplant experiment; **A)** Mean adult body size (mg of DM) of control and warming populations kept in their ancestral and novel environment; **B)** Mean somatic growth rate (per day) of control and warming populations kept in their ancestral and novel environment; the bars represent standard error of the mean.

Mean growth rate showed no significant treatment effect (LMM, df = 1; F = 0.05; P = 0.82) and no environment × treatment interaction (LMM; df = 1; F = 0.96; P = 0.33) (Figure 9B).

## 5. Discussion

Our multigenerational experiment demonstrates that gradual thermal deterioration reshapes life-history strategies in *Daphnia magna* in ways that differ from the patterns observed under abrupt temperature shifts. The most striking pattern was a distinct, rate-dependent trajectory, with an initial phase of elevated reproduction followed by the loss of the reproductive advantage at a sub-lethal fecundity threshold well below lethal temperatures (typically 35 °C for *D. magna*; Seefeldt & Ebert, 2019). This is consistent with other studies showing that demographic tipping points can occur before lethal thermal limits are reached (Kingsolver & Huey, 2008; Seefeldt & Ebert, 2019). Populations exposed to sustained warming matured earlier, grew faster, and initially produced more offspring than controls, but also exhibited shorter lifespan, and the eventual loss of reproductive advantage relative to control near 30 °C, well below their lethal limit. These findings highlight that the rate of environmental change influences both the trajectory and limits of adaptive responses, with implications for predicting persistence under ongoing climate change.

### Warming maintains fecundity until a threshold is reached

In contrast to many abrupt-shift experiments that report uniformly reduced fecundity of *D. magna* under elevated temperatures (for example, Cambronero et al., 2018; Wang et al., 2023), warming lines in our experiment maintained a fecundity advantage for most of the thermal trajectory. This pattern is consistent with studies showing that moderate warming can boost reproductive output via accelerated maturation and shorter generation times (for example, Musolin et al., 2010). However, this advantage diminishes near 30 °C, a tipping point also indicated by our thermal tolerance assay, where only 28% of individuals reproduced successfully and reproduction nearly ceased at 33 °C. The timing of extinctions in the main experiment closely paralleled this threshold, with 75% of warming populations lost between 30 and 32 °C. These results show that reproductive ceilings can be reached well before mortality thresholds, reducing population viability during warm periods even if temperatures remain below lethal limits.

### Adult body size decline was unaffected by warming

We detected an overall reduction in mean adult body size, but without difference between warming and control populations. A plausible explanation is that the temperature window that our warming populations experienced (the 26 - 30°C range) lies near the upper end of the thermal performance curve for *Daphnia* (Seefeldt and Ebert, 2019; Wang et al., 2023), where body size plasticity may already be constrained. Furthermore, there is evidence that resource limitations become stronger at higher temperatures and larger body size, causing growth to decelerate at a smaller size at warmer temperatures (Hoefnagel et al., 2018).

### The role of clutch interval

Some traits, such as clutch interval, showed limited average change but exhibited significant genotype-by-treatment interactions: the Tr clone reduced its clutch interval under warming, a shift not seen in the other clones. Because shorter intervals between clutches can substantially increase lifetime reproductive output in *Daphnia* (Garbutt et al., 2014; Claska & Gilbert, 1998), this clone-specific change may represent an important adaptive pathway under warming.

### Warming reduces lifespan

Elevated temperature significantly reduced the lifespan of experimental populations relative to control populations. Moreover, we observed a significant effect of clonal identity, with An clone showing higher longevity relative to the other two clones. The negative correlation between temperature and longevity has previously been reported (for example, Musolin et al., 2007; Ciota et al, 2014; Giovaninni et al., 2023), including the studies with *D. magna* (for example, MacArthur and Baillie, 1929; Hoefnagel et al., 2018). Our study refined these estimates by providing detailed lifespan data across a narrower but ecologically critical thermal window (26 - 30°C), capturing a sharp decline relative to control populations during the first three assays (26 - 28°C), followed by the equalization of mean longevity between the two treatment groups at the two highest temperatures.

### Life-history traits correlations

Despite an overall reduction in mean adult body size in both treatment groups, there is evidence that a larger body size was advantageous at elevated temperatures through its effect on fecundity. Moreover, we detected a significant plastic response in both traits, manifested through a simultaneous increase of both mean body size and fecundity of Tr clone when subjected to elevated temperature. Warming populations showed both a significant decline of development time and an acceleration of somatic growth rate relative to controls, consistent with findings in other studies (Henning-Lucass et al., 2016; Hoefnagel et al., 2018). These two traits showed a strong negative correlation, driven by divergent clone responses. We also saw a significant positive correlation between development time and adult body size, and a significant negative correlation between growth rate and body size. Together, these patterns suggest that delayed maturation can be advantageous at high temperatures by allowing individuals to reach a larger size coupled with a higher fecundity, whereas faster-growing clones (namely, An) show trade-off between both size and fecundity with increased rate of development. By contrast, we detected no covariation between fecundity and lifespan, nor between lifespan and development or growth, suggesting that lifespan reductions were driven primarily by direct thermal stress rather than by indirect trade-offs (Sun et al., 2023; Schwartz et al., 2016).

### Clone-specific resilience

Marked differences among clones underline the role of standing genetic variation. The Tr clone maintained larger body size and higher fecundity under warming. In contrast, the An clone was characterized by both reduced mean body size and fecundity but had greater longevity. These patterns echo mesocosm and laboratory findings showing that genotypes vary in their thermal sensitivity and capacity for persistence under stress (Geerts et al., 2015; Henning-Lucass et al., 2016). Such variation indicates the potential for evolutionary rescue (Gomulkiewicz and Holt, 1995) but also means that population-level means can mask the presence of more resilient genotypes.

### Plasticity versus genetic change

The reciprocal transplant experiment (RTE) revealed a trait-specific pattern that was not apparent in the main assays. While mean adult body size declined similarly across treatments during the main experiment, warming-line offspring were consistently smaller than controls in the transplant assay, regardless of rearing environment. This suggests a heritable component or persistent trans-generational effect in body size reduction, which was masked in the main experiment by parallel size declines across both treatments. By contrast, growth rate showed no significant treatment or environment interaction, indicating that it remained largely plastic within the time frame of the study. Notably, control offspring exhibited an increase in growth rate when reared in the novel (warming) environment, whereas warming-line offspring did not show a corresponding change when returned to the cooler environment. This trait-specific pattern aligns with other Daphnia studies showing that some traits respond primarily through evolution, others through plasticity (Yampolsky et al., 2014; Seefeldt & Ebert, 2019).

More broadly, our main experiment and RTE combined highlight that gradual warming can reveal distinct evolutionary trajectories across traits: some (like adult body size) showing persistent lineage-level shifts, while others (like growth rate) remain flexible but not divergent.

The limited number of generations (25) prior to the transplant likely constrained the magnitude of genetic change, but the divergence we detected suggests that even under relatively short time spans, gradual thermal deterioration can produce heritable shifts in key life-history traits.

### Rate of change as a key driver

The +1 °C per 4–6 generations regime produced trajectories distinct from those under single-step warming: an initial period of reproductive advantage, which was lost in sublethal temperature. This dynamic mirrors microbial experiments where gradual change extended persistence without preventing eventual decline (for example, Killeen et al., 2017). Our results reinforce that warming rate and not just absolute temperature should be treated as a primary parameter in both experiments and models, as predicted by eco-evolutionary theory and observed in microbial systems (Collins and de Meaux, 2009; Bell and Gonzalez, 2011; Killeen et al., 2017).

## Limitations

Our experiment used a limited number of genotypes and spanned only 25 generations before the transplant assay. Longer-term experiments incorporating greater genetic diversity are needed to fully capture the interplay between plasticity and evolution under gradual warming. In addition, our design did not manipulate resources or community composition, which may influence whether the fecundity advantage observed here emerges across a wider range of ecological contexts.

Although some baseline differences were present (e.g., slightly higher fecundity in Tr under warming and longer lifespan in controls at 26 °C), these did not alter the main trajectories: warming populations maintained a fecundity advantage until the highest temperature and consistently showed lifespan declines. This suggests that the observed patterns were robust to initial conditions.

## Broader implications and conclusions

From an applied perspective, our findings show that gradual warming can mask an impending demographic decline by temporarily enhancing reproduction. Sub-lethal thresholds such as the ∼30 °C fecundity ceiling may be crossed more often in the future as thermal safety margins shrink, particularly in regions or seasons already near species’ optima (Deutsch et al., 2008). In *D. magna*, this tipping point occurred near 30 °C, well below the lethal limit, and was closely aligned with extinction events. The fact that reproductive ceilings can be reached long before mortality thresholds, and that their timing depends on the pace of warming, underscores the need to treat warming rate as a fundamental driver in both experimental design and predictive modelling, and to recognise that identical thermal endpoints reached at different rates can produce qualitatively different demographic and evolutionary outcomes (Killeen et al., 2017). As thermal safety margins shrink globally, such rate-dependent demographic ceiling may become a common feature of population decline, making the detection of early-warning signals for reproductive tipping points essential for anticipating and mitigating biodiversity loss under climate change.

## 6. Appendix

### 6.1 Outlier detection results and model selection

#### Span between the clutches

Two extreme values were identified as potential outliers using Grubbs’ test (p < 0.05) and were excluded from model comparison.

#### Neonate body size

Four extreme values were identified as potential outliers using Grubbs’ test (p < 0.05) and were excluded from model. Model selection was based on a comparison of AIC values and a likelihood ratio test. A simplified model excluding interaction terms (AIC = −1274.3) was favored over a more complex model that included the interactions (AIC = −1263.1), as the later did not significantly improve model fit according to a likelihood ratio test (χ^2^, P = 0.9).

#### Adult body size

A single extreme value was identified as potential outliers using Grubbs’ test (p < 0.05) and was excluded from model.

#### The development time

Two extreme values were identified as potential outliers using Grubbs’ test (p < 0.05) and were excluded from model comparison. A simplified model excluding interaction terms (AIC = 1198.7) was favored over the full interaction model (AIC = 1208.5; LRT P = 0.77).

#### Somatic growth rate

Model selection was based on a comparison of AIC values and a likelihood ratio test. A simplified model excluding interaction terms (AIC = - 107.35) was favored over a more complex model that included the interactions (AIC = -97.86), as the later did not significantly improve model fit according to a likelihood ratio test (χ^2^, P = 0.72).

### 6.2 Non-significant correlations between life history traits

We found no significant correlation between lifespan and development time (r = - 0.04; t (79) = - 0.34; P = 0.73); lifespan and somatic growth rate (r = 0.02; t (33) = 0.09; P = 0.93); lifespan and fecundity (r = 0.06; t (69) = 0.52; P = 0.61); fecundity and span between clutches (r = - 0.28; t (272) = -0.29; P = 0.78); adult body size and span between clutches (r = - 0.03; t (67) = - 0.26; P = 0.79); somatic growth rate and fecundity (r = - 0.08; t (60) = - 0.6; P = 0.55).

## Acknowledgements

This study has been funded by Tübitak (project code: TB.0064).

We are grateful to the former postgraduate student, Seray Altıntaş, for assistance in development of the protocol for algal quantification, and for her assistance with pilot experiments and maintenance of cultures. We thank the undergraduate students, including Helin Bahar Calı, Kerem Durdu, Gülnur Karabiber, and Ece Türksever, for their assistance with pilot experiments and maintenance of cultures. We are grateful to Dr Bebac Šmepčević for great illustrations within Figure and for assistance in development of the protocol for algal quantification.

## Data Accessibility Statement

Data is available through the Dryad Digital Repository (http://datadryad.org/share/LINK_NOT_FOR_PUBLICATION/ZkFUNcfZSiCaQRD6FLZIJFfQ5TEIUAQLMD3KHbT3ars).

